# The interacting effects of climate and land-use/land-cover changes on ecological communities

**DOI:** 10.1101/2023.12.07.570587

**Authors:** Kimberly L. Thompson, Jonathan M. Chase, Ruben Remelgado, Carsten Meyer

**Author notes:** Correspondence to: Kimberly L. Thompson.

## Abstract

Human activities continue to create land-use/land-cover (LULC) change across the Earth’s surface, and together with climate change, are major drivers of changes in biodiversity through time. However, the impacts of these spatiotemporally variable drivers on biodiversity change can be complex. We examined the effects of interactions between climate and LULC change on bird communities across the continental United States over nearly three decades. We analyzed temperature and precipitation data alongside data on tree-canopy, cropland, urban, and surface-water cover to understand how climate/LULC-change interactions influence species richness and abundance. Our results revealed stable or increasing trends in species richness, but a decline in overall abundance primarily from common species and a consequent increase in aspects of evenness of communities. We found that areas experiencing warming and drying climates exhibited increased species richness and slower declines in abundance. However, impacts of LULC change had contrasting effects on richness and abundance. Areas that experienced increasing tree-canopy cover over time had increasing trends in species richness, but exacerbated declines in abundance. On the other hand, areas with increasing cropland had moderated abundance declines, but more declining trends in richness. Finally, we found that the effects of climate/LULC-change interactions varied across the range of each pressure. While some interactions support a dominant role of climate change in structuring communities, others indicate that LULC change can mitigate or exacerbate the impact of climate change on biodiversity. Overall, our results highlight the importance of considering the direction and magnitude of each driver when assessing how climate and LULC interactions shape ecological communities.

## Introduction

Human activities exert numerous, simultaneous pressures on Earth’s biodiversity, which have resulted in global declines in both species richness and abundance of many taxa (Diaz *et al*., 2019). Notably, however, local and regional trends of biodiversity change are often incongruent with these global declines (Dornelas *et al*., 2023, Dornelas *et al*., 2014, Vellend *et al*., 2013). Instead, there are many instances where local biodiversity stays the same or even increases, and these can counteract the instances where local biodiversity is declining when multiple studies are compared (Blowes *et al*., 2019, Dornelas *et al*., 2023, Dornelas *et al*., 2019).

Central to understanding differences in the trends of biodiversity at local scales is a concurrent understanding of the anthropogenic pressures to which ecological communities are exposed (Gonzalez *et al*., 2023, Oliver & Morecroft, 2014). Not only do individual pressures, such as land-use change or climate change, vary spatiotemporally in their intensity (Ji *et al*., 2014, Kuemmerle *et al*., 2016), so also do the ways in which different pressures interact. For example, fragmentation of forest habitats can lead to decreases in land surface temperatures, but the magnitude of these declines varies from tropical to boreal ecosystems (Mendes & Prevedello, 2020).

Land-use/land-cover (LULC) change and climate change represent two of the most dominant pressures on biodiversity (IPBES, 2019). LULC change is responsible for the majority of contemporary extinctions (Caro *et al*., 2022, Maxwell *et al*., 2016), while climate change has forced local extirpations and range shifts of many taxa in many places (Antao *et al*., 2022, Urban, 2015). Since communities typically experience LULC and climate change simultaneously, the interactions between these pressures (and similarly those between other global change pressures) are especially critical for our understanding of biodiversity change (Newbold, 2018, Oliver & Morecroft, 2014, Peters *et al*., 2019).

As the most abundant group of terrestrial vertebrates (Lees *et al*., 2022), birds are an ideal taxon for deepening our understanding of how interactions between climate and LULC change impact ecological communities. Since birds can move freely between habitats, they are strong indicators of ecosystem health (Rosenberg *et al*., 2019, Thorn *et al*., 2018) and can play an important role in the functioning of ecosystems (e.g. pollination (Zanata *et al*., 2017), seed dispersal (Sekercioglu *et al*., 2004), trophic regulation (Böhm *et al*., 2011)). Furthermore, long-term data on entire bird communities are often available over broad spatial extents owing to their easy observability and interest to scientists and lay people alike (Tulloch *et al*., 2013), which can facilitate our understanding of how biodiversity change resulting from climate and LULC interactions extends from local communities to continents.

Importantly, bird communities are also highly sensitive to both climate and LULC (Ferger *et al*., 2017, Haddou *et al*., 2022). In terms of climate, aspects of both temperature (e.g., Howard *et al*., 2020, Peters *et al*., 2016) and precipitation (e.g., McCain, 2009, McCain & Colwell, 2011) strongly influence patterns of bird richness and abundance. For LULC, conversion from natural to anthropogenic land uses (e.g., agricultural lands, urban areas) also impacts bird communities (Allen *et al*., 2019, Gregory *et al*., 2007, Heldbjerg *et al*., 2018). While change in temperature is often more influential than LULC for bird communities (Ferger *et al*., 2017, Howard *et al*., 2020, Saunders *et al*., 2022, Spooner *et al*., 2018), our understanding of how these two major pressures interact in the face of their inherent spatiotemporal variability to drive richness and abundance changes in ecological communities remains limited (Newbold *et al*., 2019).

Expectations of how climate and LULC change interact to impact biodiversity tend more towards negative synergies (i.e., the combined impact is more detrimental than the impact of either individual pressure) (Brook *et al*., 2008, Newbold *et al*., 2019) due to the ways in which LULC change can prevent species from tracking their climate niches (Frishkoff *et al*., 2016). For example, habitat fragmentation due to agricultural expansion or urban sprawl can impede dispersal and potentially subject species to climate conditions outside of their optimal range (Brook *et al*., 2008, Opdam & Wascher, 2004). Under this scenario, we might expect (eventual) declines in both richness and abundance as species and individuals become unable to persist in their communities (Haddou *et al*., 2022).

However, interactions between climate and LULC change can either augment or dampen biodiversity change depending on the particular combination of pressures and species-specific resource requirements (Fong *et al*., 2018). For example, Frishkoff et al. (2016) found that bird species affiliated with drier climates had a higher probability of occupying areas converted to agriculture than those that favored wetter environments. Consequently, even if an area experiences agricultural conversion, richness and abundance could increase if colonization by species better suited to the emerging environmental conditions exceeds extirpation.

In forested areas, microclimates resulting from heterogeneity in land cover create climate-change refugia since they do not track broader climatic changes (Oliver & Morecroft, 2014), and can therefore support higher numbers of individuals and species (Stein *et al*., 2014). If, however, communities have species that are highly sensitive to a particular aspect of climate or LULC (e.g., a species with a specific habitat requirement or a narrow thermal range), the impact of either pressure is likely to override the effect of the other (Viana & Chase, 2022). For example, in an evaluation of the relationship between rates of climate warming, rates of LULC change, and changes in abundance in terrestrial birds, Spooner et al. (2018) found that populations declined when temperatures increased more rapidly; however, there was no effect of LULC change or the interaction of temperature and land-use. Since biodiversity responses to anthropogenic pressures are complex and nonuniform (Spooner *et al*., 2018, Thomas *et al*., 2006), outcomes in which climate/LULC interactions generate positive and negative biodiversity change, as well as those in which one pressure is dominant, are all plausible.

Here, we use nearly 30 years of data from a long-term survey of bird communities across the continental United States (i.e., the North American Breeding Bird Survey; Pardieck *et al*., 2020) to investigate how trends in LULC change, climate change, and interactions between these pressures have affected trends in bird richness and abundance. Specifically, we paired temperature and precipitation data with state-of-the-art data on fine-scale changes in tree-canopy, cropland, urban, and surface-water cover to address how interactions between climate and LULC change have driven changes in bird communities. Examining LULC change, climate change, and biodiversity change with this broad spatial and temporal scope allows us to better understand the conditions under which ecological communities thrive and decline in the presence of interacting anthropogenic pressures.

## Methods

### Data and Preprocessing

#### Breeding Bird Survey Data

The North American Breeding Bird Survey (BBS) dataset consists of bird point count data from thousands of roadside survey routes in the United States and Canada (Pardieck *et al*., 2020). Each route is approximately 39.2 km and is surveyed during the bird breeding season, with the majority of surveys occurring in May or June. During the surveys, expert observers stop every 800 meters (i.e., 50 stops in total) and conduct a 3-minute point count, during which they record every bird seen or heard within a 400-meter radius.

We downloaded BBS data, which also include metadata describing the survey conditions, route characteristics, and species identities (Pardieck *et al*., 2020). We additionally downloaded the coordinates for all surveyed routes (United States Geological Survey, 2020). To pair the BBS data with gridded data on LULC and climate (see Predictor Data below), we limited the study extent to the contiguous United States and included only routes that had been surveyed during the period of 1992–2018. Although climate data are available for longer periods, this period represents the range for which we were able to compile reliable, high-quality LULC data (Remelgado *et al*., 2023). We further excluded routes surveyed for less than 20 years since short time series have been shown to bias estimates of change (Leung *et al*., 2020). Finally, we removed four routes that had duplicated spatial coordinates, leaving a sample size of 1,758 routes (Figure 1).

**Figure 1:**
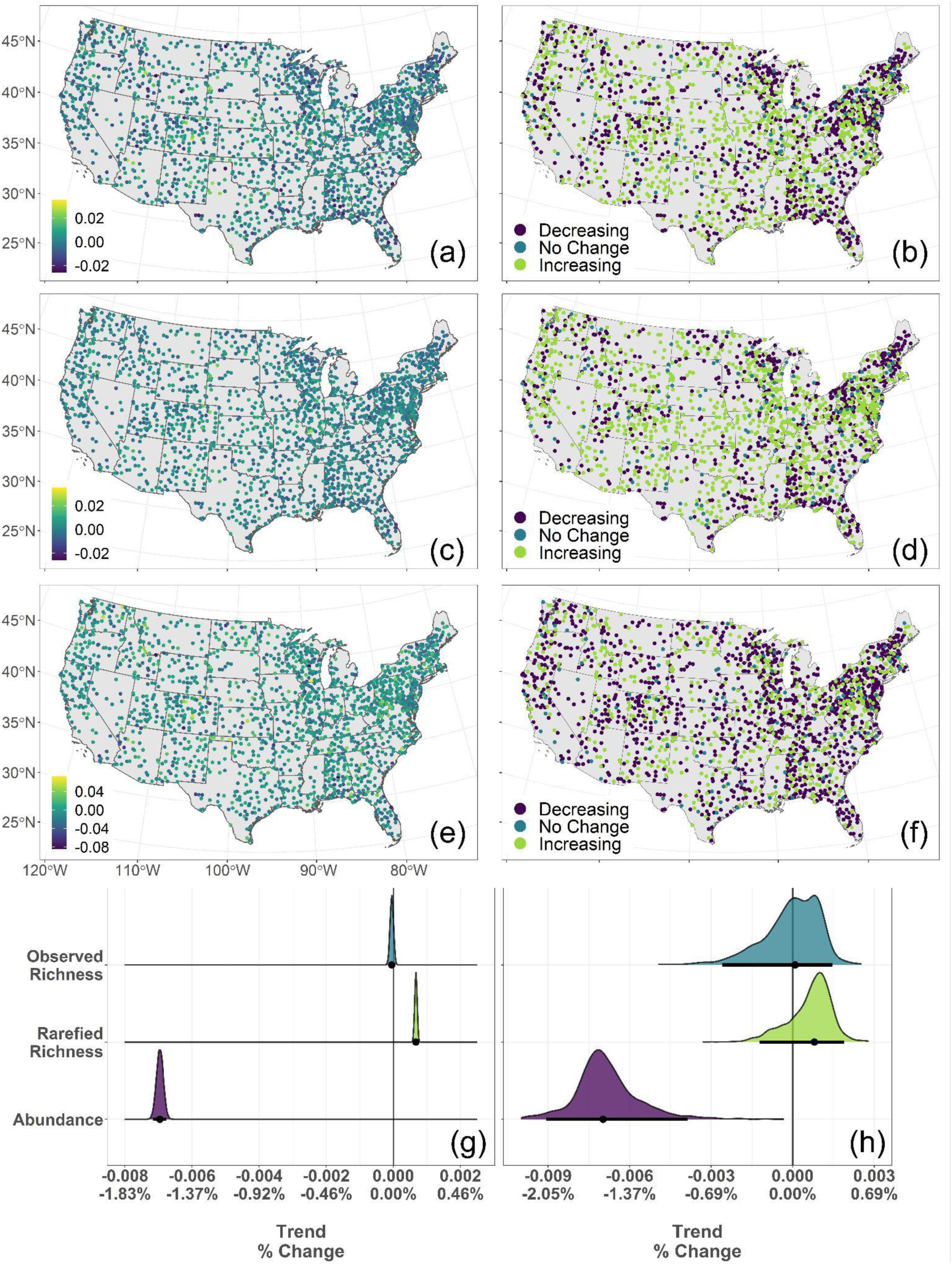
Raw and directional trends in observed species richness: (a), (b); rarefied species richness (c), (d); and abundance (e), (f) for 1,758 bird communities surveyed for at least 20 years from 1992 to 2018 as part of the North American Breeding Bird Survey. (g) Posterior distribution of the intercept, signifying the baseline conditions in 1992, and (h) posterior predictions for observed species richness, rarefied species richness, and abundance. For (g) and (h), the black point represents the mean trend value and the black line the 95% credible interval.

For each BBS route, we calculated the observed species richness and total abundance in each year. Because sampling effort was approximately equal for each route, we could directly compare trends. Furthermore, to determine how species richness was influenced by changes in the total abundance of individuals for each route and time point, we calculated individual-based rarefied richness, which compares species richness values for a given number of individuals (Gotelli & Colwell, 2001). We calculated rarefied richness by finding the minimum number of individuals observed across all routes and years (minimum = 30 individuals), multiplying this value by 2 (60 individuals), and extracting the rarefied richness value for this number of individuals for each route in each year (Chao *et al*., 2014).

### Predictor Data Climate

We acquired gridded precipitation (mm) and temperature (°C) data from the PRISM climate dataset (PRISM Climate Group, 2020), which has a resolution of 4 kilometers across the contiguous United States, for the months of May and June (i.e., the months during which the majority of bird surveys took place) in each year (1992–2018). We then calculated the two-month average in each grid cell in each year to obtain summarized values of May/June precipitation and temperature.

### Land-Use/Land-cover

#### Tree-canopy Cover

To ensure that our analyses included smaller tracts of tree-canopy cover, rather than only those officially designated as forest, we used data on continuous tree-canopy cover (Remelgado & Meyer, 2023). These data result from a multi-scale modeling framework combining different remote sensing products and exploit path-dependencies in forest change to reconstruct tree-canopy cover dynamics between 1992 and 2018 at a 300-m resolution. As a result, these data resolve historical gaps in satellite archives that impede observing annual changes in the 1990s (Remelgado *et al*., 2023).

### Cropland

We used the European Space Agency’s Climate Change Initiative (CCI) global land cover product, mapped for the period of 1992–2018 with a 300-meter resolution (European Space Agency, 2017). The original data inadvertently map implausible changes caused by the extension of its time series into the 1990s with 5-km AVHRR satellite data, compared to the 300-m data used in later years. To tackle this deficiency, we used data on cropland extents mapped in 4-year intervals since the turn of the century (Potapov *et al*., 2022) to verify the annual occurrence of this land use. Using the verified pixels, we then mapped per-pixel percentages of cropland at a 300-meter resolution, aggregating the following classes: ‘rainfed cropland’, ‘irrigated or post-flooding cropland’, and ‘mosaic cropland greater than 50% with natural vegetation less than 50%’ (i.e., CCI land cover classes 10, 20, 30).

### Urban Areas

As a proxy for changes in the degree of urbanization, we used the GAIA (Global Artificial Impervious Areas) dataset (Gong *et al*., 2020), which maps the first year of impervious cover between 1985 and 2018 at a 30-meter resolution. We split these data annually to depict annual extents of impervious cover and truncated the time series to 1992–2018. Then, we aggregated the data into a 300-m resolution grid through averaging. This resulted in an annual time series of per-pixel percentages of impervious cover. Taken together, these percentages of impervious cover represent varying degrees of development (i.e. ranging from roads and highways to villages and cities). Hereafter, we use any change in the amount of impervious cover as a proxy for whether an area is becoming more (or less) urban.

### Permanent and Seasonal Water

We used annual data on surface-water seasonality from the Global Surface Water dataset (Pekel *et al*., 2016), which was mapped between 1984 and 2021 at a 30-meter resolution. For each pixel, these data distinguish water as either ‘seasonal’ or ‘permanent’. Permanent water refers to pixels that were classified as water-covered in every satellite image (i.e., 16 daily Landsat images). If water was not found in every image, pixels were classified as containing seasonal water cover. For this study, we focused on maximum water extents or the combined coverage of permanent and seasonal water. We aggregated these data into an annual time series with a 300-meter resolution for 1992–2018 through averaging, resulting in time series of per-pixel water cover percentages.

### Associating Predictors with BBS Routes

To determine the value of each predictor associated with each BBS route in each year, we first buffered each route by 400 meters (i.e., the distance at which observers record birds during surveys). Then, for each predictor, we calculated the weighted average of the cells that overlapped with each BBS route’s buffered area, with the weights determined by the proportion of the buffered area within a cell. This resulted in a single value of each predictor for each BBS route in each year. **Analysis**

### Trend Calculation

To facilitate comparisons between each biodiversity metric, we log-transformed observed richness, rarefied richness, and abundance values. We then determined the trend of each log-transformed biodiversity metric and each predictor (mean May/June temperature (°C), mean May/June precipitation (mm), tree-canopy cover (%), cropland cover (%), urban cover (%), and water cover (%)) with generalized linear models that employed an autoregressive process of order one. For each variable and route combination, we calculated the trend as the slope of the fitted line, along with its standard error (i.e., the uncertainty in the trend). To facilitate visualization of the biodiversity trends across space, we standardized slopes by their standard errors and then classified them as ‘increasing’, ‘decreasing’, or ‘no change’ by constructing histograms with 50 bins for each response. We defined data points that fell within the bins immediately adjacent to zero as ‘no change’, while those above and below were classified as ‘increasing’ and ‘decreasing’, respectively (Figure 1).

### Modeling

For trends in log-transformed observed richness, log-transformed rarefied richness, and log-transformed abundance, we created separate Bayesian regression models with Gaussian error distributions. We accounted for uncertainty in the modeled biodiversity trend values by including the standard errors of these trends as part of the response. Predictors were the same for all models and included standardized trends in temperature, precipitation, tree-canopy cover, cropland, urban cover, and water cover, as well as two-way interactions between climate and LULC change variables (e.g., trends in temperature and tree-canopy cover, temperature and cropland, precipitation and tree-canopy cover, precipitation and cropland, etc.).

For all models, we used default priors for the intercept, but weakly informative priors for predictors, informed by estimated trends of climate and LULC change in the United States (Supplementary Methods). We ran all models for 10,000 iterations with 1,000 warmups over 4 chains and examined model convergence using Rhat values (Gelman-Rubin diagnostic) and visually through trace and density plots (Gelman & Rubin, 1992).

Prior to modeling, we evaluated the spatial autocorrelation of each biodiversity response by calculating Moran’s I using MCMC sampling with 999 iterations, but found non-significant values for all responses (Figure S1). After modeling, we calculated the model residuals for each iteration/chain combination (9,000 iterations * 4 chains) and found the mean residual for each route (i.e., sample point). We again tested for spatial autocorrelation by calculating Moran’s I for the model residuals and found non-significant values (Figure S1).

We examined the sensitivity of our results to biases in surface-water cover data, which can arise due to improvements in Landsat coverage that occurred during our study period and consequently overestimate the amount of increasing surface water (Remelgado *et al*., 2023). For each BBS route, we applied the Mann-Kendall test to time series of both surface-water coverage and Landsat data quality, represented by the number of months in a year with data. Since the Mann-Kendall test is a non-parametric analysis that identifies significantly increasing or decreasing trends, we were able to identify routes that had significant trends in both the surface-water and data-quality time series, indicating that for these routes, the changes in surface-water cover may have been driven by changes in Landsat data quality. Out of the 1,758 sites in our sample, we identified 57 routes that had potentially biased trends in surface-water cover. To test whether the inclusion of these routes affected our results, we reran each Bayesian model excluding those 57 routes, and compared the parameter estimates for surface-water cover change. Since the parameter estimates and credible intervals for surface-water change were nearly the same between each model (Table S1), we proceeded with the full sample of 1,758 routes.

Finally, we also examined the sensitivity of our results to uncertainty in predictor trends by rerunning each model using the trend values of the predictors plus or minus each predictor trend’s standard error.

All analyses, as well as all pre-processing procedures, were conducted using R Statistical Software (v4.2.2; R Core Team, 2022). We used the mobr package to calculate individual-based rarefied richness (McGlinn *et al*., 2021), the prism, terra, raster, and sf packages for extraction and processing of the climate and LULC data (Hart & Bell, 2015, Hijmans, 2021a, Hijmans, 2021b, Pebesma, 2018), the nlme package for trend calculations (Pinheiro et al. 2020), the brms package for modeling (Bürkner, 2017), and the ggplot2, tidybayes, ggdist, and viridis packages for visualizations (Garnier *et al*., 2021, Kay, 2021, Kay, 2022, Wickham, 2016).

## Results

### Trends in Biodiversity, Climate, and Land cover/use

Overall, we found that the mean trends in observed and rarefied richness were positive, but very close to zero (Observed Richness: [-0.025, 0.034], µ=0.0002; Rarefied Richness: [-0.017, 0.026], µ=0.001, Figure 1a, 1c, S2). Relative to richness measures, abundance trends were more variable, but the overall mean was negative (Abundance: [-0.084, 0.072], µ=-0.006, Figure 1e, S2). Trends for all biodiversity measures showed minimal spatial clustering with opposite trends often observed at adjacent locations (Figure 1b, 1d, 1f).

Average trends in temperature and precipitation were both positive (Temperature: [-0.035, 0.102], µ=0.043, Precipitation: [-2.628, 4.420], µ=0.564, Figure S2, S3). While warming and wetting trends occurred throughout the entire United States throughout the study period, the highest magnitude trends were concentrated in the central and eastern regions (Figure S3). Although there were very few BBS routes that experienced cooling trends (Figure S2, S3), many routes experienced drying, particularly those in the western United States (Figure S2, S3).

Tree-canopy cover trends generally indicated losses, and these were strongest in the southeastern United States (Figure S2, S3), with annual losses in these areas between 0.75% and 1.5% (Tree-canopy: [-1.415, 0.0643], µ=-0.136, Figure S2, S3). Cropland was mostly stable (Figure S2), though we registered small gains throughout the US, with the exception of the southwest, which had relatively little cropland cover from the start (Cropland: [-0.007, 0.003], µ=-0.0001, Figure S3). Urban areas either remained unchanged or increased through time (Urban: [0.000, 1.597], µ=0.064, Figure S2, S3). These increases were distributed spatially, but occurred predominantly in the central and eastern US, and along the western coast. Finally, water cover generally increased, with the majority of routes experiencing at least small gains (Seasonal and Permanent Water: [-0.279, 0.829], µ=0.0731, Figure S2, S3).

### Baseline and Overall Change

When climate and LULC predictors did not change (i.e., trend values = 0), we found no change in observed richness (µ=-0.01% per year, 95% credible interval -0.03–0.01%), small increases in rarefied richness (µ=0.15% per year, 95% credible interval 0.14–0.17%), and declines in abundance (µ=-1.59% per year, 95% credible interval -1.63– -1.54%) (Figure 1g). For the entire period of 1992–2018, we found no net change in either observed or rarefied richness (observed richness: µ=-0.01% per year, 95% credible interval -0.60–0.34%; rarefied richness: µ=0.15% per year, 95% credible interval -0.28–0.46%), but declines in abundance (µ=-1.60% per year, 95% credible interval -2.36–-0.90%) (Figure 1h). Despite these overall estimates, density distributions for each biodiversity metric contained routes that were changing both positively and negatively, with abundance having the greatest spread (Figure S2a–c).

### Impact of Climate and LULC on Trends of Species Richness

We found that areas with warming temperatures and stable or increasing tree-canopy cover experienced gains in species richness, while areas with cooling temperatures and declining tree-canopy cover showed losses of species richness (Figure 2a–c). In areas that were becoming colder, increases in tree-canopy cover were associated with no change in observed species richness (i.e., credible intervals for the different levels of tree-canopy cover change overlapped zero), while both stable and decreasing tree-canopy cover in cooling areas resulted in observed species losses (Figure 3a). Patterns differed between observed and rarefied richness, however. For rarefied richness, tree-canopy cover had little to no impact in areas that were getting colder (Figure 3b). Additionally, while warming trends benefitted observed species richness more or less equally for all levels of tree-canopy cover change (i.e., credible intervals for each level of tree-canopy change overlapped one another, Figure 3a), rarefied richness had the highest increases where warming and increased tree-canopy cover occurred simultaneously (Figure 3b).

**Figure 2:**
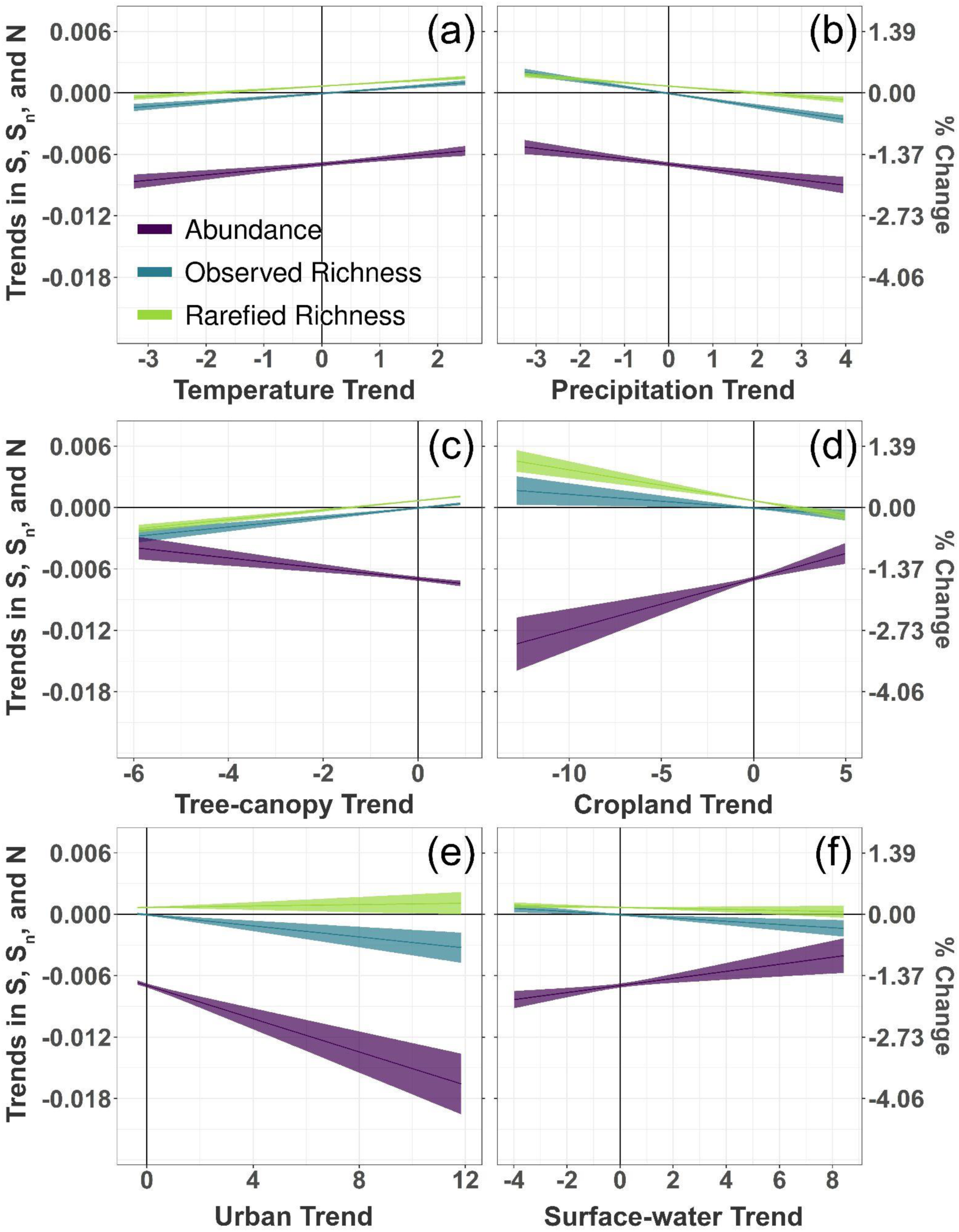
Conditional effects of trends in temperature (a), precipitation (b), tree-canopy cover (c), cropland cover (d), urban cover (e), and surface water cover (f) on trends in log-transformed observed richness, rarefied richness, and abundance generated from Bayesian regression models. Models were built with biodiversity, climate, and land-use trend data for 1,758 bird communities surveyed for at least 20 years from 1992 to 2018 as part of the North American Breeding Bird Survey. Uncertainty bars represent 95% credible intervals.

**Figure 3:**
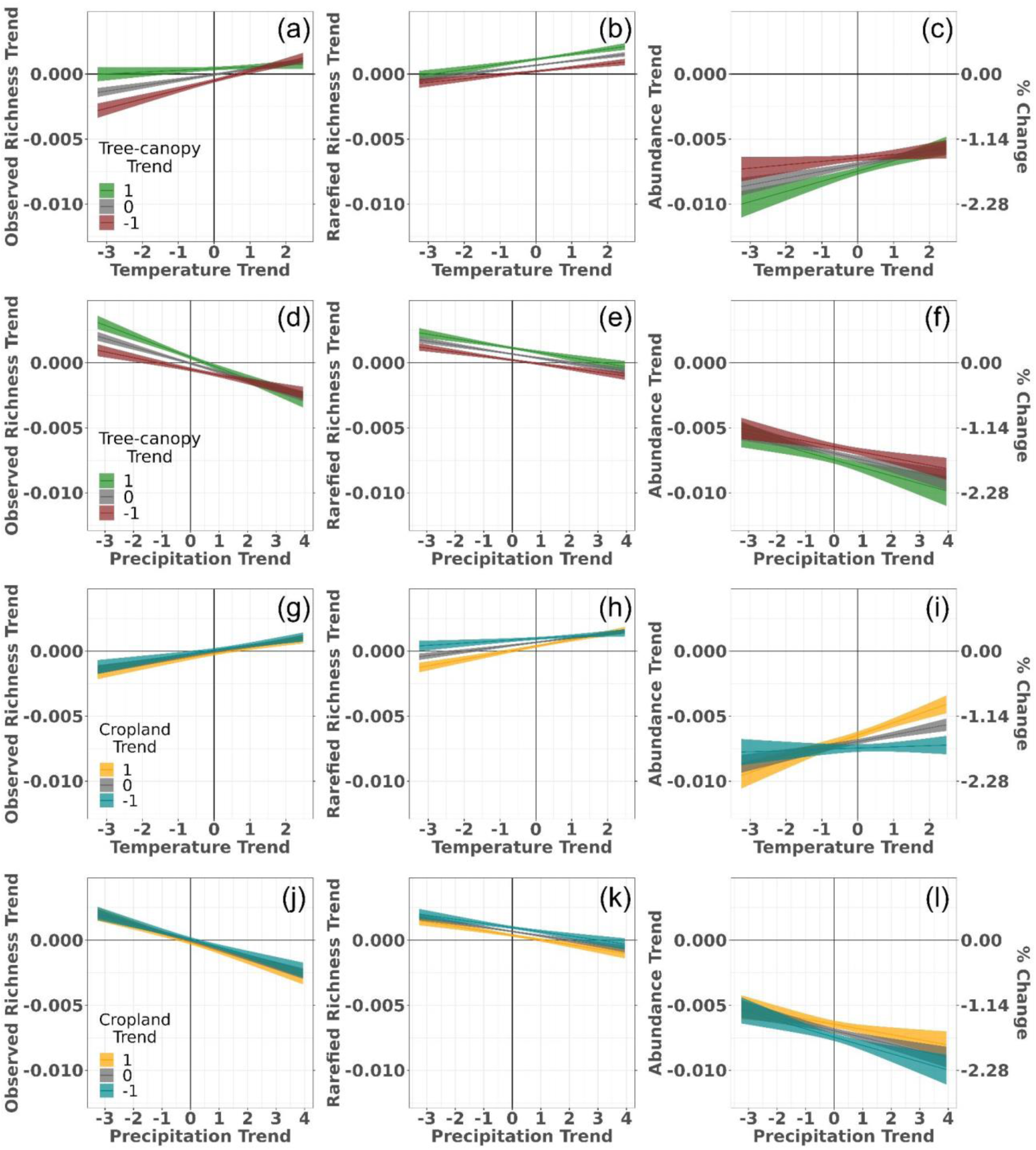
Conditional effects of the interactions between changes in temperature and tree-canopy cover (a–c), precipitation and tree-canopy cover (d–f), temperature and cropland (g–i), and precipitation and cropland (j–l) on biodiversity trends (log values) in observed species richness, rarefied species richness, and abundance generated from Bayesian regression models. Models were built with biodiversity, climate, and land-use trend data for 1,758 bird communities surveyed for at least 20 years from 1992 to 2018 as part of the North American Breeding Bird Survey. Uncertainty bars represent 95% credible intervals.

Areas that became drier tended to show increases in species richness, while species richness tended to decline in areas becoming wetter through time (Figure 2b). Richness declines with increases in precipitation were stronger for observed richness than for rarefied richness, while species gains due to drying were consistent between the two responses (Figure 2b). The effect of reduced precipitation varied across levels of tree-canopy cover, with more species richness gains occurring in areas that were becoming drier and also increasing in tree-canopy cover than in drying areas with unchanging or declining tree-canopy cover (Figure 3d, 3e).

Declines in cropland cover were associated with gains in both observed and rarefied richness, while increasing cropland cover resulted in modest species declines (Figure 2d). Despite this, there was no difference in observed species richness across the different levels of cropland change when interacting with temperature, with cooling areas showing declines and warming areas showing increases (Figure 3g). In contrast, the interaction of cropland change and cooling temperatures resulted in rarefied richness changes ranging from small decreases to no change when cropland cover was stable, and no change to small increases when cropland cover decreased through time (Figure 3h).

Similar to the results for temperature, we found that changes in precipitation through time had a stronger influence on richness trends than changes in cropland through time. Regardless of how cropland changed through time, gains in observed species richness tended to occur in areas drying through time, whereas losses in observed species richness occurred in areas becoming wetter through time (Figures 3j). This effect was consistent for rarefied richness, except that species declines were not as severe in areas that were becoming wetter (Figure 3k).

Generally, both increasing urban cover and increasing water cover had no impact on rarefied richness trends, but were associated with declines in observed richness trends (Figure 2e, 2f). Changes in urban cover were less influential than temperature and precipitation changes, with observed and rarefied richness gains occurring in areas that were becoming warmer and drier, regardless of changes in urban cover (Figure 4a–b, 4d–e). Shifts in surface-water cover interacted with temperature change; both observed and rarefied richness losses were reduced in areas that were cooling and that also had stable or declining surface water coverage (Figure 4g, 4h). Alternatively, areas that were becoming cooler and also had increases in their surface-water coverage had more severe declines in species richness (Figure 4g, 4h). For rarefied richness, we also found that areas warming through time had relatively more species richness gains when there was also increasing surface-water coverage (Figure 4h). Finally, we found that areas that were increasing in precipitation and surface-water coverage had the highest observed species losses (Figure 4j), while decreases in rarefied richness were not as strong (Figure 4k).

**Figure 4:**
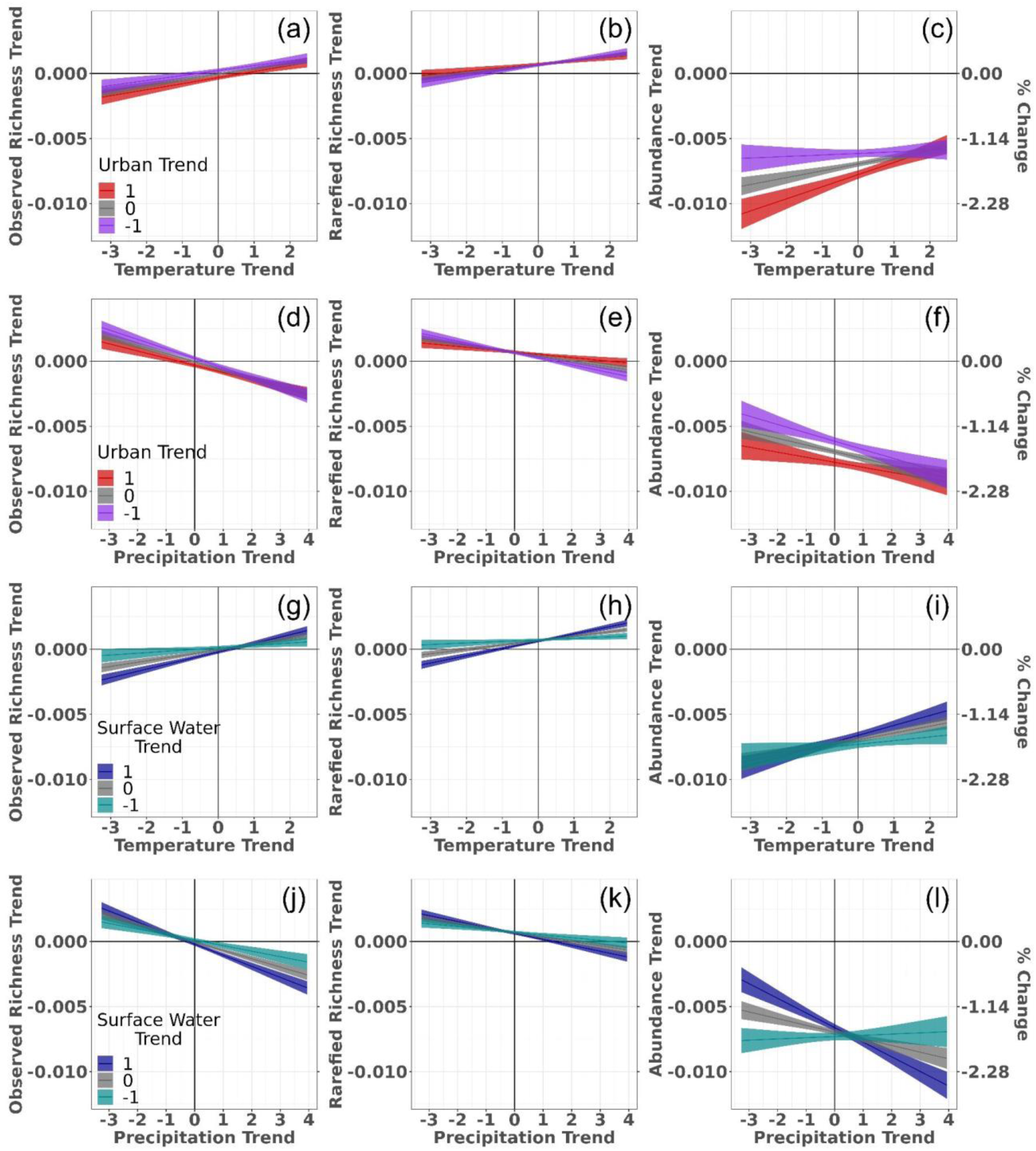
Conditional effects of the interactions between changes in temperature and urban cover (a–c), precipitation and urban cover (d–f), temperature and surface water cover (g–i), and precipitation and surface water cover (j–l) on biodiversity trends (log values) in observed species richness, rarefied species richness, and abundance generated from Bayesian regression models. Models were built with biodiversity, climate, and land-use trend data for 1,758 bird communities surveyed for at least 20 years from 1992 to 2018 as part of the North American Breeding Bird Survey. Uncertainty bars represent 95% credible intervals.

### Impact of Climate and LULC on Trends of Abundance

The total abundance of birds within each route tended to decline across the entire range of all climate and LULC change predictors; however, both climate and LULC influenced the magnitude of these declines. For example, there were relatively fewer individuals lost in areas that were becoming warmer and experiencing declines in tree-canopy cover (Figure 2a, 2c). However, warming temperature trends were more influential than tree-canopy cover trends, as the impact of warmer temperatures did not differ between different levels of tree-canopy cover change (Figure 3c). On the other hand, fewer individuals were lost in areas that were becoming cooler and also declining in tree-canopy cover than in areas that were becoming cooler and also increasing in tree-canopy cover (Figure 3c).

Fewer individuals were also lost in areas where cropland cover was increasing (Figure 2d), but temperature seemed to override this effect in places that were becoming colder, since abundance losses due to cooling did not differ across levels of cropland change (Figure 3i). Nonetheless, in areas that were getting warmer, those that were simultaneously experiencing an increase in cropland cover lost relatively fewer individuals compared to areas with stable or declining cropland (Figure 3i).

For precipitation, fewer individuals were lost in places that were becoming drier (Figure 2). Neither tree-canopy change nor cropland change impacted changes in abundance due to drying (Figure 3b). While losses of individuals still occurred in places that were becoming drier, not only were they less than losses found in areas that were becoming wetter, but there was no difference between different levels of tree-canopy cover or cropland change (Figures 3f, 3l).

Increasing urban cover led to strong declines in abundance (Figure 2e), which were exacerbated by interactions with temperature (Figure 4c). For areas experiencing cooling temperatures, urbanizing areas experienced the highest magnitude declines in abundance, but as temperatures warmed, these declines stabilized for all levels of urban-cover change (Figure 4c). Contrary to the interaction with temperature, urban-cover change impacted abundance trends only when precipitation levels were stable or slightly changing (i.e., slightly increasing or decreasing) (Figure 4f). At these small magnitude precipitation changes, more individuals were lost in urbanizing areas, while at larger magnitude precipitation changes, losses of individuals were not distinguishable based on urban-cover change (Figure 4f).

Increases in the coverage of surface water reduced declines in abundance (Figure 2f); however, when interacting with temperature, the different levels of surface-water cover change were associated with abundance declines of similar magnitude (Figure 4i). More abundance declines occurred in cooler areas and fewer declines occurred in warming areas regardless of the change in surface-water coverage (Figure 4i). For areas experiencing drying, increasing surface-water coverage coincided with a reduction in the loss of individuals such that abundance declines were dramatically lower in areas that were increasing their surface-water coverage (Figure 4l). This effect shifted for areas that were getting wetter, with simultaneous increases in surface water associated with stronger declines in abundance than for areas with stable or decreasing surface-water coverage (Figure 4l).

For all models, using the upper and lower uncertainty values for the trends of predictors did not change the overall pattern of the results; we refer the reader to the supplement for figures corresponding to main text figures 2–4 for models that used the upper and lower standard errors of the predictors (Figures S4–S9).

## Discussion

We used detailed data on changes in land use/land cover (LULC) (including tree-canopy, cropland, urban, and surface-water cover) and climate (temperature and precipitation) to examine how interactions between LULC and climate change have impacted bird richness and abundance in the continental United States over the last 30 years. Our results revealed stable or increasing species richness, but a decline in abundance, indicating a loss of individuals from common species and increased evenness in communities. We also found evidence of opposing dynamics between richness and abundance due to their differing responses to tree-canopy cover and cropland change. Overall, our results show that different interactions between climate and LULC matter in different ways for richness and abundance, highlighting that the relationship between global change pressures and biodiversity is complex and nuanced (see Dornelas *et al*., 2023 for a more comprehensive review of this issue). Consequently, a one-size-fits-all solution to mitigate biodiversity change is unlikely to be successful.

### The Nature of Community Change

Trends in species richness and abundance were both impacted by trends in climate and LULC; however, richness responses were variable, while abundances tended to decline across the entire range of climate and LULC change variables. These abundance declines are consistent with previous estimates that approximately three billion individual birds have been lost in North America since 1970 (Rosenberg *et al*., 2019). Similarly, the variable responses in species richness coincide with evidence regarding the diverse influence of climate and LULC on richness patterns, as some species are highly influenced by these factors, while others are less affected (Hobi *et al*., 2021).

Despite widespread declines in abundance, species richness and abundance both responded favorably to warming temperatures and declining precipitation. Rising temperatures due to climate change can be detrimental to biodiversity (Nunez *et al*., 2019); however, we found that increasing temperatures—up to a change of 2.5°C and in the absence of other changes in precipitation or LULC—generally led to higher species richness and reduced declines in abundance. A study that examined the risk of extirpation among montane bird species under future climate change scenarios came to similar conclusions and projected that increases in temperature over the next 100 years would lead to only a 3% increase in extirpation risk (McCain & Colwell, 2011). While warmer temperatures may confer energetic savings for some species (e.g. shortened incubation periods (Bleu *et al*., 2017), reduced energetic stress due to warmer winters (Zuckerberg *et al*., 2011)), the prevailing expectations for birds suggest that positive responses to warming temperatures would occur predominantly in communities in cooler environments, whereas communities in warmer climates would exhibit negative responses (Sumasgutner *et al*., 2023). However, our results demonstrate richness increases and less intense declines in abundance in areas in both warm and cool climates throughout the United States, highlighting that positive biodiversity responses to increasing temperatures can occur irrespective of climatic zone (Figure 1, S3) (Peel *et al*., 2007).

Our observations of the positive response of species richness and the less negative response of abundance to drier conditions were unexpected since wetter environments typically support more resources for birds (Liu *et al*., 2013). At the same time, intense precipitation events may restrict foraging times and are consequently associated with reduced survival and fecundity (Gullett *et al*., 2015). Correia et al. (2015) found that both richness and abundance were dramatically lower in warmer and drier areas along the Iberian peninsula, possibly due to the lower tree cover and reduced resource availability in these areas rather than climate per se. Accordingly, understanding the mechanisms behind community responses to changes in precipitation may only be possible while simultaneously considering LULC changes.

In response to LULC changes, we found opposing patterns between species richness and abundance. Increasing tree-canopy cover promoted species richness but exacerbated declines in abundance. Alternatively, increasing cropland mitigated declines in abundance, but triggered declines in richness. The conflicting responses between richness and abundance in these two land-cover types highlight that global change pressures can influence different dimensions of biodiversity in different ways. Relative to cropland, tree-canopy cover is often associated with higher levels of environmental heterogeneity (e.g., due to vertical structuring) and therefore can provide more niche space for multiple species to coexist (MacArthur & MacArthur, 1961, Tews *et al*., 2004). Cropland, on the contrary, is often characterized by monocultures that lack numerous and diverse niches. At the same time, agricultural areas offer predictable and abundant food resources, which could explain why abundance declines were less severe when cropland increased. Regardless of the mechanisms, the divergent responses of richness and abundance trends to changes in tree-canopy cover and cropland demonstrate the challenge of generalizing biodiversity responses and the potential choices that can emerge in efforts to conserve ecological communities.

Due to the mathematical linkages between different measures of biodiversity (Jost, 2010, Shimadzu, 2018), similarities (or differences) in the direction of observed richness, rarefied richness, and abundance trends can shed light on how and why communities are changing (Blowes *et al*., 2022). For example, because rarefied richness controls for changes in abundance while remaining sensitive to changes in relative abundance (Gotelli & Colwell, 2001), a positive relationship between observed richness and abundance, with rarefied richness more or less unchanged, indicates community change resulting primarily from shifts in overall abundance. Alternatively, cases that deviate from this pattern indicate that community change does not result from shifts in overall abundance alone, but from shifts in the relative abundance of species (Jost, 2010). Consequently, the declines in abundance, together with the simultaneous increases or stability in richness that we found imply that the relative abundance and evenness of communities changed throughout the time series (Blowes *et al*., 2022). Specifically, we expect that communities likely lost individuals from more common species. These results differ from the frequent expectation that biodiversity loss tends to be concentrated on rarer species more than common ones (i.e., resulting in decreased evenness) (Gaston, 1994) due to the high sensitivity of rare species to both LULC change (Newbold *et al*., 2018, Newbold *et al*., 2013) and climate change (Foden *et al*., 2013, Pearson *et al*., 2014). However, our results, as well as other recent work (Gregory *et al*., 2023, Rigal *et al*., 2023), show that declines in the abundance of common bird species, leading to increased evenness, may sometimes be prevalent.

The relationship between observed and rarefied richness can also reveal critical information about the nature of community change. For example, a positive relationship between observed and rarefied richness indicates that pressures affecting the relative abundance of species, rather than the overall abundance of communities, are predominant (Blowes *et al*., 2022).

Generally, observed and rarefied richness trends had the same sign and direction in response to trends in climate and LULC, reinforcing the idea that these anthropogenic pressures have disproportionate impacts on species (Beissinger *et al*., 2023). Notably, richness responses to urbanization did not follow this pattern, with trends in rarefied richness increasing slightly while trends in observed richness declined. This suggests that urbanization affects biodiversity primarily via changes in the overall abundance of communities, which makes sense given the reduced and degraded habitats that result from this type of LULC change (Marzluff, 2001). Additionally, since our results revealed universal abundance declines in response to trends in climate and LULC pressures, we provide further evidence that shifts in the overall abundance of communities and the relative abundances of species within these communities can occur simultaneously to elicit biodiversity change (Allen *et al*., 2019, Rosenberg *et al*., 2019).

### The Nature of Climate-change/LULC-change Interactions

Previous work on interactions between climate and LULC in bird communities has generally found that climate, particularly temperature, exerts a stronger influence on biodiversity trends than LULC (Ferger *et al*., 2017, Howard *et al*., 2020, Saunders *et al*., 2022, Spooner *et al*., 2018). While our results confirmed the dominance of climate in certain interactions (i.e., the credible intervals of positive (+1) and negative (-1) LULC change overlapped the credible interval indicating zero LULC change, sensu Côté *et al*., 2016), we found that whether climate change overrode the impact of LULC change depended on the magnitude and direction of the climate change, the magnitude and type of LULC change, and the metric used to detect biodiversity change. Consequently, even in interactions where temperature and precipitation change superseded the impact of LULC change, this dominance was not consistent across the range of potential climatic changes, the range of potential LULC changes, or across the different measures of biodiversity. The one exception was the interaction between precipitation and cropland, where wetter conditions were associated with negative biodiversity outcomes and drier conditions were associated with positive biodiversity outcomes for both species richness and abundance regardless of changes in cropland. That neither species richness nor abundance responded to changes in cropland cover when there were simultaneous changes in precipitation is surprising since other research has shown that agriculture can exacerbate the negative effects of increased precipitation on fledging success (Garrett *et al*., 2022) and impact differences in occupancy between wet and dry areas (Frishkoff *et al*., 2016).

While there are several potential mechanisms underlying how climate-/LUCC-change interactions affect ecological communities, the examples in which the dominance of climate corresponded with biodiversity declines suggest the potential for physiological stress (Brown *et al*., 2004, Navarrete *et al*., 2021, Speakman & Król, 2010) and reduced resource availability (Socolar *et al*., 2017) induced by temperature and precipitation changes. This could in turn lead to changes in habitat suitability and the necessity to emigrate to more favorable areas (Heinrichs *et al*., 2016). Alternatively, the instances in which climate change was more influential than LULC change and gave rise to an increase (or reduced decline) in biodiversity imply the opposite: energetic savings, increased resource availability, and improved habitat suitability (Hawkins *et al*., 2003, Schaefer *et al*., 2008, Sumasgutner *et al*., 2023). However, the levels of LULC change were relatively low compared to the levels of temperature and precipitation change. Therefore, it is possible that climate change may not be the dominant force behind biodiversity change in areas with higher magnitude LULC changes. At the same time, because climate change is expected to intensify in the coming decades, areas with stable temperature and precipitation are likely to become less prevalent (IPCC, 2023) and may override the impacts of LULC change in more places in the near future.

We also found cases in which temperature or precipitation change did not override the effects of LULC changes on biodiversity change (i.e., the credible intervals of positive (+1) and negative (-1) LULC change did not overlap the credible interval indicating zero LULC change). These instances demonstrate the capacity of LULC change to augment or dampen the richness and abundance changes expected due to climate change alone. In fact, despite evidence for the primary role of temperature and precipitation in structuring bird communities (Howard *et al*., 2020), all of the LULC types we tested mitigated (or exacerbated) the effects of climate change across part of the range of climatic changes and for at least one of the biodiversity metrics.

Overall, this highlights how critical interactions between climate and LULC change are for understanding the complexities of biodiversity change.

### The Geography of Community Change

Over the near-30-year period of our study, we did not uncover any definitive regional patterns of richness and abundance change (Figure 1). Instead, we found only small clusters of local communities (routes) that displayed similar trends in richness and/or abundance. This heterogeneity across regional and continental scales emphasizes the importance of local environmental conditions in structuring ecological communities. While approximate regional and continental patterns emerged in some of the climate and LULC pressures (e.g., temperature, precipitation, tree-canopy cover) (Figure S3), there was also considerable heterogeneity in the geographic patterns of the different global change pressures. Consequently, since our results demonstrate that the effects of climate/LULC interactions on bird communities vary according to the direction and magnitude of each pressure, the geographic heterogeneity of community trends therefore seems inevitable.

## Conclusion

We show that the interplay between climate and LULC change has profound implications for biodiversity dynamics of bird communities over a near-30-year time series in the United States. However, this interplay is highly variable across a range of climatic changes, LULC changes, and biodiversity measures. Although the positive responses to warming temperatures, drying conditions, and certain LULC changes hint at the adaptive potential of ecological communities, diverse outcomes are likely and we need to account for this breadth in our strategies to conserve biodiversity. The distinct responses of species richness and abundance to certain LULC changes, the variation in interactions where climate change was dominant, and the geographic heterogeneity of biodiversity change challenge the notion of a singular, standardized approach to mitigating biodiversity loss. Instead, targeted interventions that account for lack of uniformity in the effects of interacting global change pressures are critical.

## Acknowledgments

We thank the extended ‘Biodiversity Synthesis’ and ‘Macroecology and Society’ groups at iDiv, as well as S. Blowes and B. Rosenbaum, who provided advice on the analyses and their presentation. All authors were supported by the German Centre for Integrative Biodiversity Research (iDiv) Halle-Jena-Leipzig, funded by the German Research Foundation (FZT 118, 202548816)

