## Supplemental Methods, Figures, and Tables for "The interacting effects of climate and land-use/land-cover changes on ecological communities"

### Supplemental Materials

### Supplemental methods

*Weakly Informative Priors*

We used existing information on the rate of change of climate and land-use to define weakly informative priors for the Bayesian regression models. The IPCC reports that temperature has increased at approximately 0.2°C per decade, though likely between 0.1°C and 0.3°C (Allen *et al.*, 2018). Therefore, we assumed a normally distributed prior for the trend of temperature with a mean of 0.02°C/year and a standard deviation of 0.01°C/year.

Climate change has also intensified the water cycle leading to more intense rainfall and flooding, as well as more intense drought (Allen *et al.*, 2018). Despite these potentially offsetting extremes, precipitation has likely increased since 1950, with a faster rate of increase beginning in the 1980s. In the United States, most areas have experienced a 10% increase in annual mean precipitation (Arias *et al.*, 2021). Therefore, we assumed a normally distributed prior with a mean of 1.1 mm/year and a standard deviation of 0.5 mm/year to account for potentially high variability between regions. We did expect, though, that increased frequency of flooding and droughts would affect the coverage of water (specifically of seasonal water), but since these phenomena potentially nullify each other, we assumed a normally distributed prior for permanent and seasonal water, with a mean of 0 and a standard deviation of 1.

Although it is possible to have tree-canopy cover that does not comprise a forest, we used rates of forest change as a proxy for tree-canopy cover change. Trends of forest loss were variable, with losses of roughly 0.2–0.3 million hectares per year during 1990–2000 across North and Central America, gains of about 0.2 million hectares per year during 2000–2010, and finally losses of about 0.1 million hectares per year from 2010–2020 (FAO, 2022). In the United States specifically, forest cover declined by about 0.03% annually from 2010–2020 (FAO, 2022). Due to the presence of both gains and losses since 1990, we assumed a normally distributed prior with a mean of 0 and a standard deviation of 0.3.

The majority of the United States experienced stable cropland cover from 2000–2019, though there were some small areas in which cropland expanded by about 15% during this time period (Potapov *et al.*, 2022). Since expanding by 15% over a 20-year period would yield a rate of 0.75% per year, we assumed a normally distributed prior for cropland with a mean of 0 and a standard deviation of 0.75.

We used an interactive graph depicting the share of national populations living in urban areas over the past 500 years to approximate the rate of expansion of urban areas (Ritchie & Roser, 2018). In 1992 approximately 76.1% of the U.S. population lived in urban areas, while in 2016 that percentage increased to 86.9%, yielding a population change of about 0.23% per year. To account for urban areas expanding outward or simply becoming denser to accommodate the additional population, we assumed a low overall rate of change and defined a normally distributed prior with a mean of 0.25 and a standard deviation of 0.25.

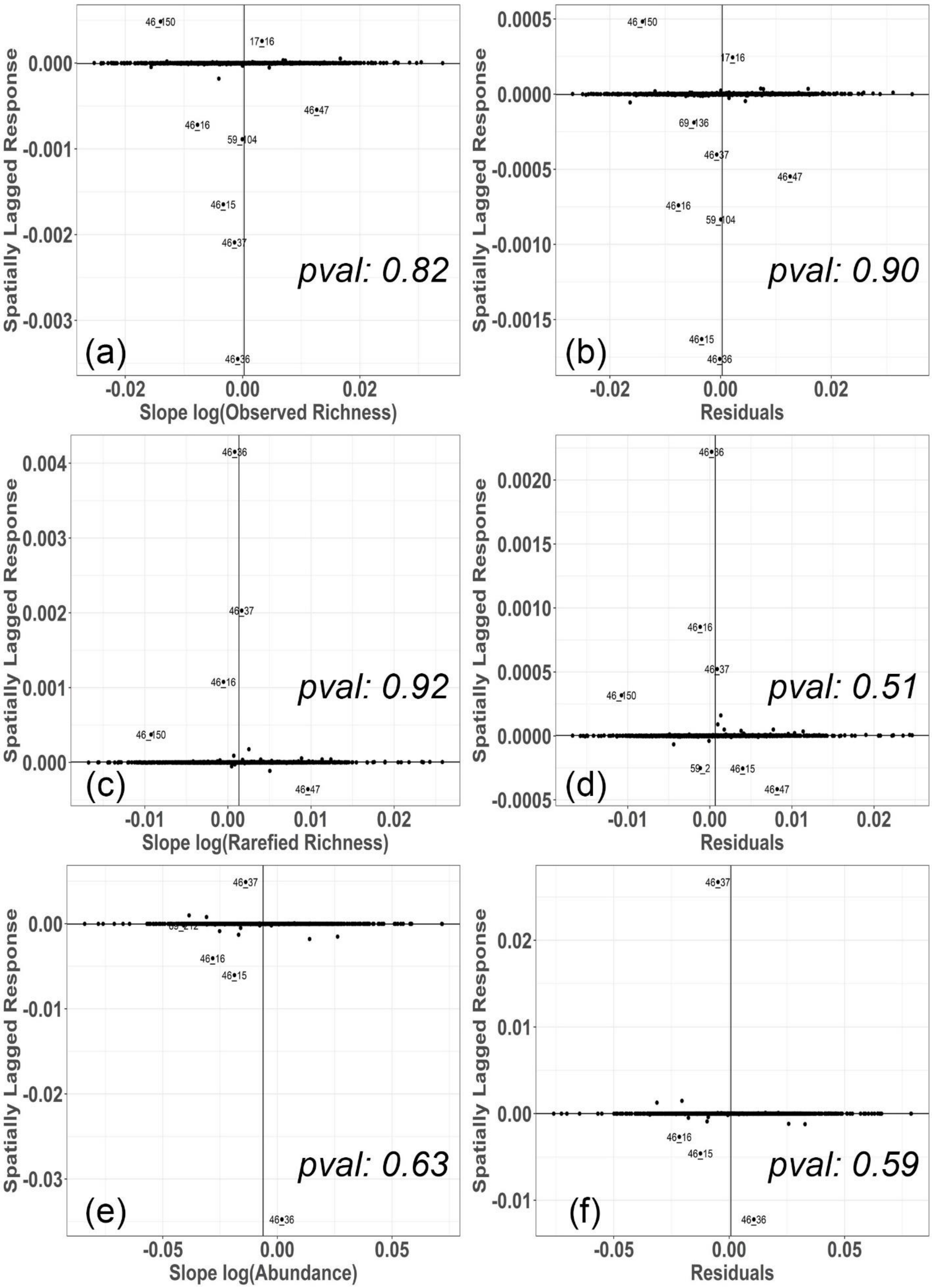

Spatially Lagged Response
Slope log(Observed Richness)
pval: 0.82
(a)
46_150
17_16
46_47
46_16
59_104
46_15
46_37
46_36
Residuals
pval: 0.90
(b)
69_136
Slope log(Rarefied Richness)
pval: 0.92
(c)
pval: 0.51
(d)
59_2
Slope log(Abundance)
pval: 0.63
(e)
69_212
pval: 0.59
(f)

Supplemental Figure 1: Moran's I scatterplots and corresponding p-values calculated for trends in the log values of observed species richness (a), rarefied species richness (c), and abundance (e), and the residuals from Bayesian regression models with the log values of observed richness (b), rarefied richness (d), and abundance (f) as the responses. Labeled on each graph represent the ID of Breeding Bird Survey routes that deviated from the zero line (i.e., no spatial autocorrelation).

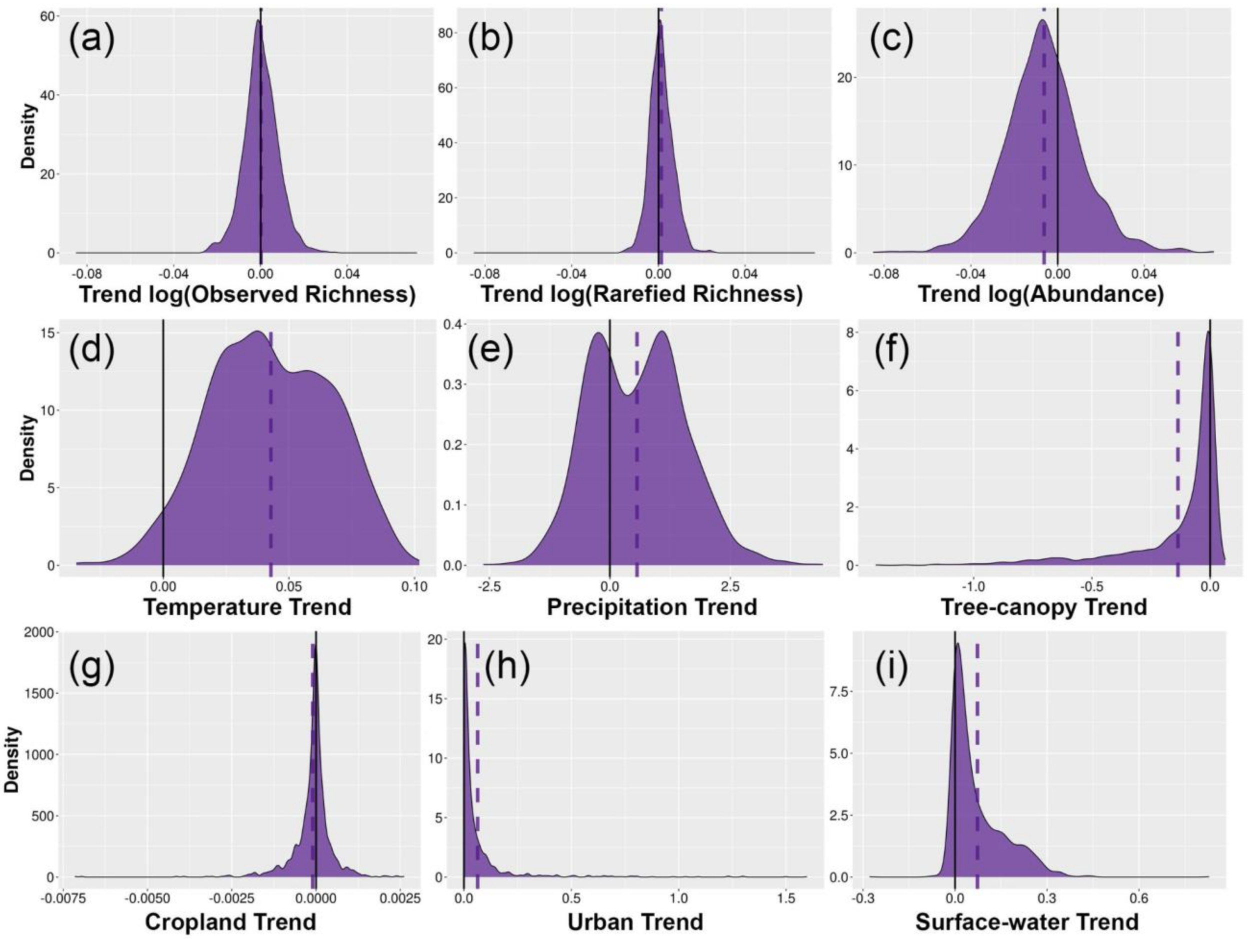


Supplementary Figure 2: Density plots showing the distribution of trends in log(observed richness) (a), log(rarefied richness) (b), log(abundance) (c), temperature (d), precipitation (e), tree-canopy cover (f), cropland (g), urban cover (h), and surface-water cover (i).

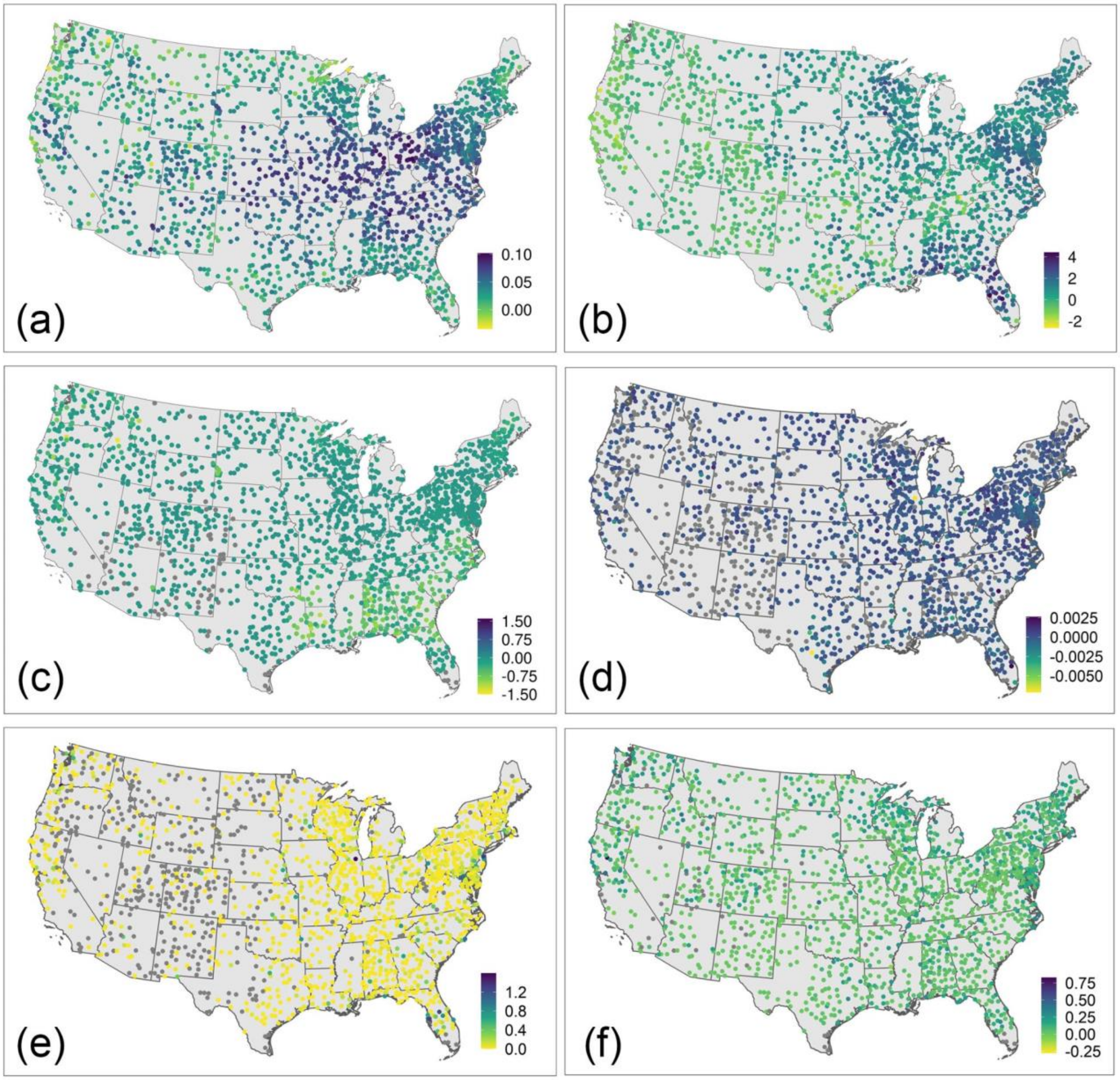


Supplemental Figure 3: Trends in temperature (a), precipitation (b), tree-canopy cover (c), cropland (d), urban cover (e), and surface-water cover (f) for 1,758 avian communities surveyed for at least 20 years within the period of 1992–2018 as part of the North American Breeding Bird Survey.

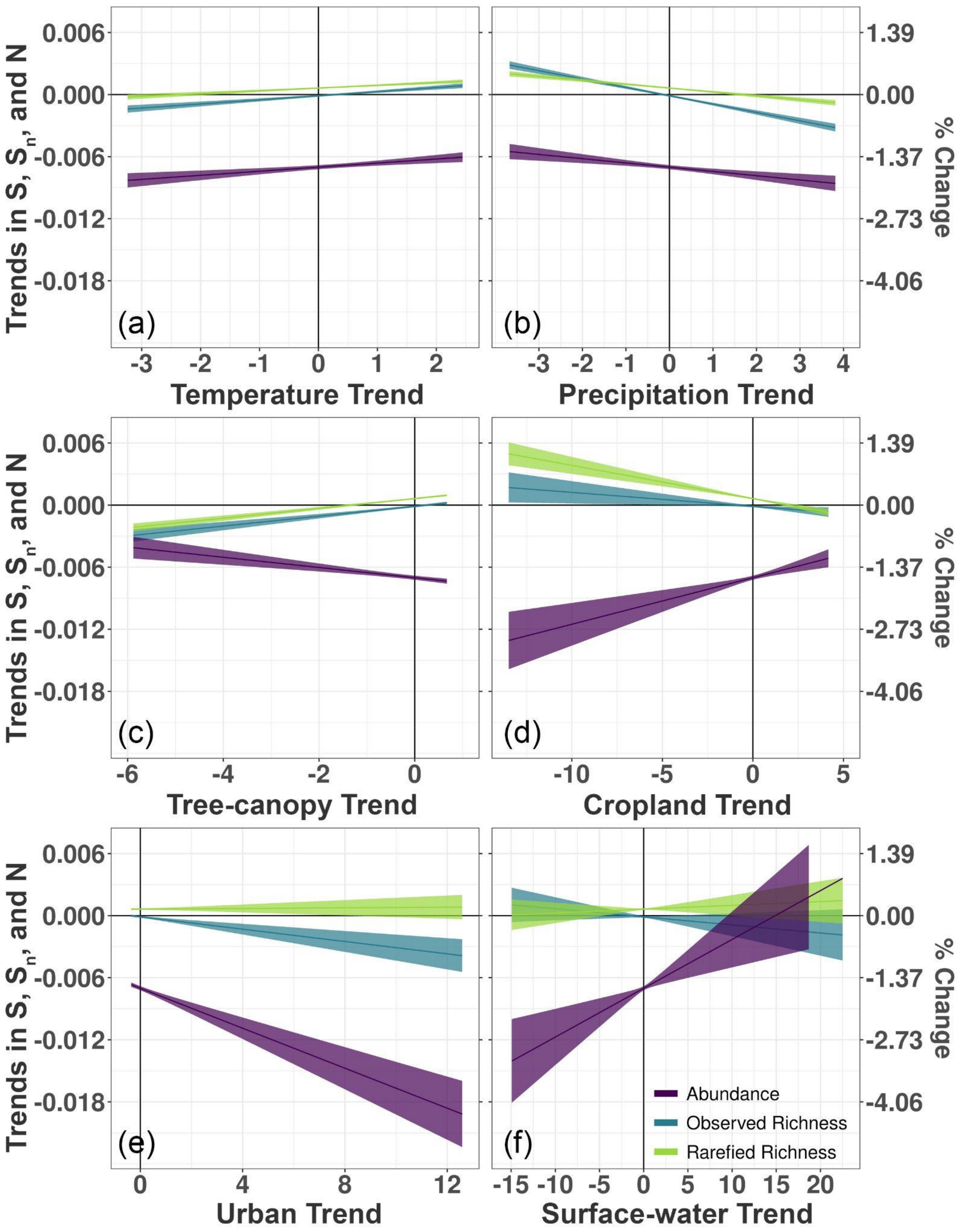
Trends in $S$, $S_n$, and N
% Change
0.006
0.000
-0.006
-0.012
-0.018
1.39
0.00
-1.37
-2.73
-4.06
(a)
(b)
(c)
(d)
(e)
(f)
Temperature Trend
Precipitation Trend
Tree-canopy Trend
Cropland Trend
Urban Trend
Surface-water Trend
Abundance
Observed Richness
Rarefied Richness

Figure S4: Lower standard error of predictors model: Conditional effects of trends in temperature (a), precipitation (b), tree-canopy cover (c), cropland cover (d), urban cover (e), and surface-water cover (f) on trends in log-transformed observed richness, rarefied richness, and abundance generated from Bayesian regression models. Models were built with biodiversity, climate, and land-use trend data for 1,758 avian communities surveyed for at least 20 years within the period of 1992–2018 as part of the North American Breeding Bird Survey; however, the trend values used for the climate and land-use predictors was the slope of the fitted line minus the associated standard error. Uncertainty bars represent 95% credible intervals.

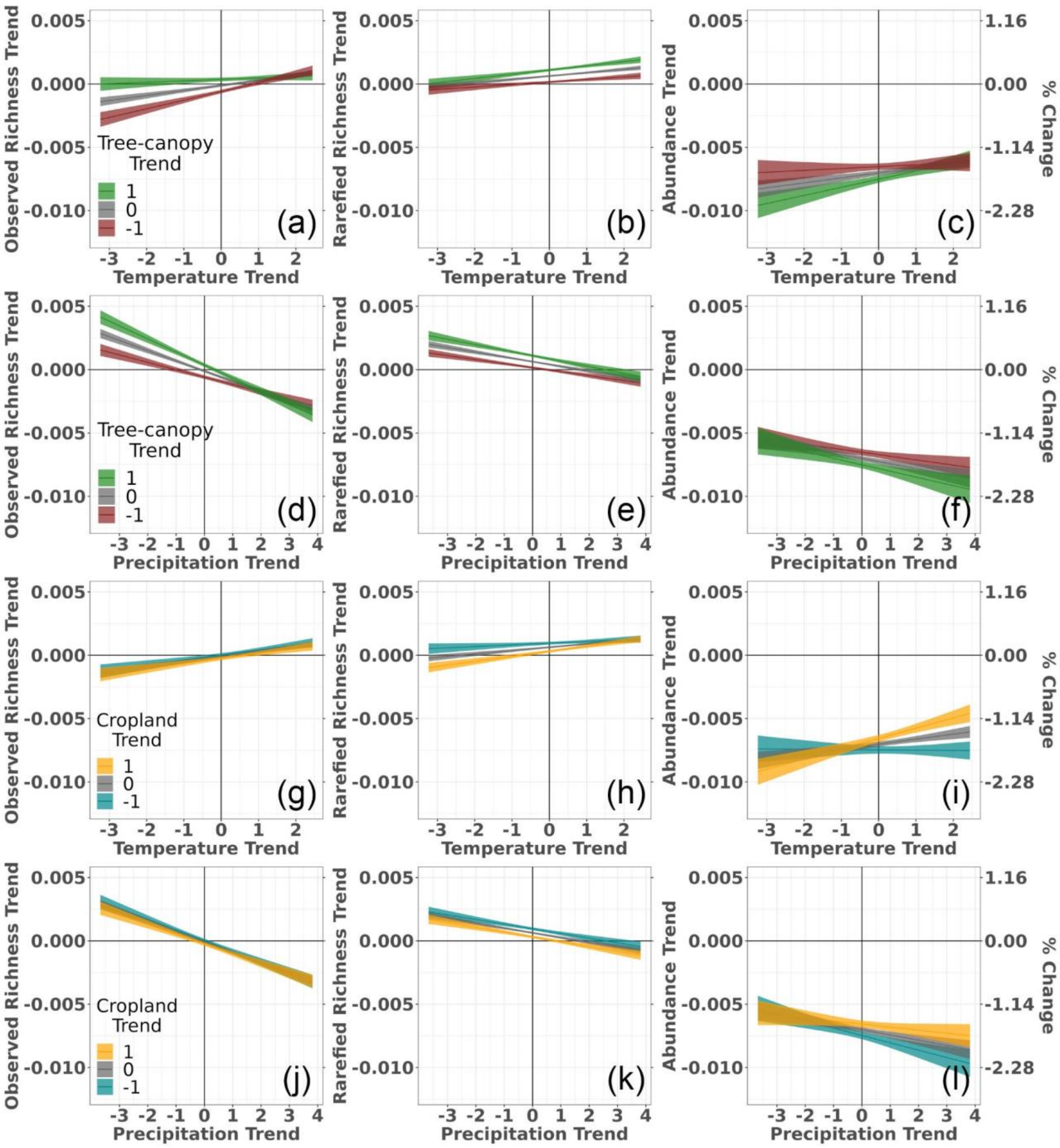


Figure S5: Lower standard error of predictors model: Conditional effects of the interactions between changes in temperature and tree-canopy cover (a–c), precipitation and tree-canopy cover (d–f), temperature and cropland (g–i), and precipitation and cropland (j–l) on biodiversity trends in log-transformed observed species richness, rarefied species richness, and abundance generated from Bayesian regression models. Models were built with biodiversity, climate, and land-use trend data for avian communities surveyed for at least 20 years within the period of 1992–2018 as part of the North American Breeding Bird Survey; however, the trend values used for the climate and land-use predictors was the slope of the fitted line minus the associated standard error. Uncertainty bars represent 95% credible intervals.

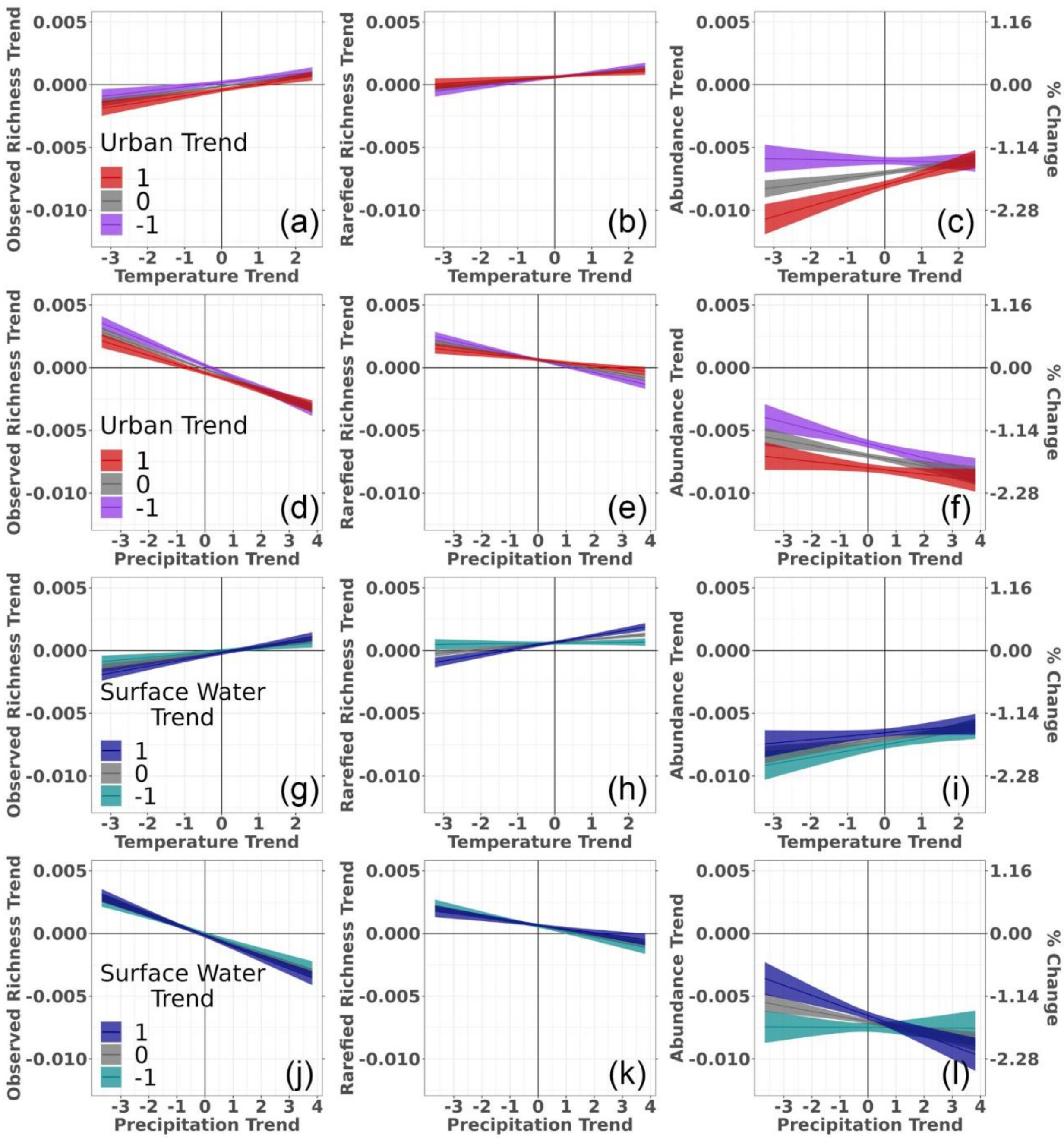


Figure S6: Lower standard error of predictors model: Conditional effects of the interactions between changes in temperature and urban cover (a–c), precipitation and urban cover (d–f), temperature and surface-water cover (g–i), and precipitation and surface-water cover (j–l) on biodiversity trends in log-transformed observed species richness, rarefied species richness, and abundance generated from Bayesian regression models. Models were built with biodiversity, climate, and land-use trend data for avian communities surveyed for at least 20 years within the period of 1992–2018 as part of the North American Breeding Bird Survey; however, the trend values used for the climate and land-use predictors was the slope of the fitted line minus the associated standard error. Uncertainty bars represent 95% credible intervals.

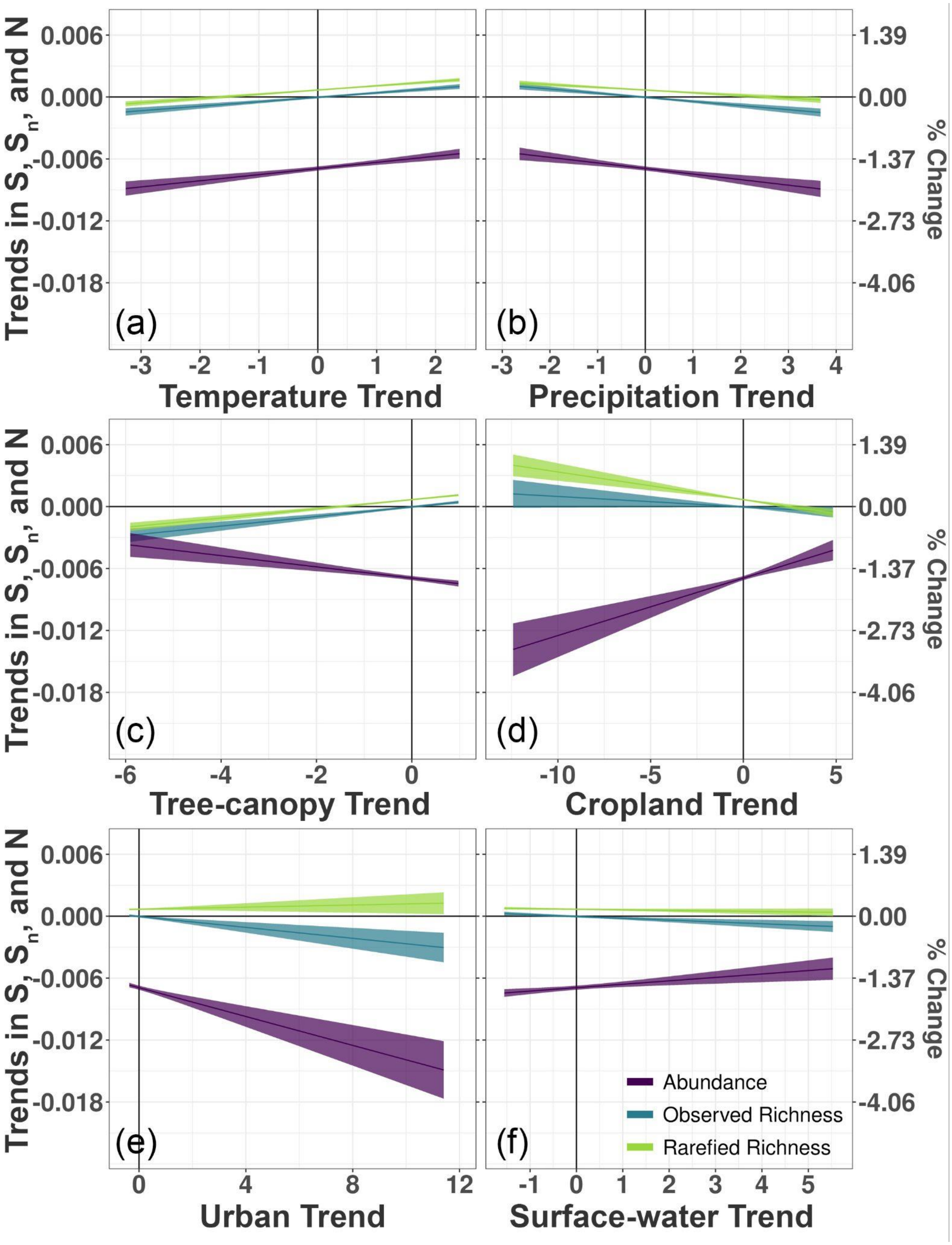

Trends in S, Sn, and N
% Change
0.006
0.000
-0.006
-0.012
-0.018
1.39
0.00
-1.37
-2.73
-4.06
(a)
(b)
(c)
(d)
(e)
(f)
Temperature Trend
Precipitation Trend
Tree-canopy Trend
Cropland Trend
Urban Trend
Surface-water Trend
Abundance
Observed Richness
Rarefied Richness

Figure S7: Upper standard error of predictors model: Conditional effects of trends in temperature (a), precipitation (b), canopy cover (c), cropland cover (d), urban cover (e), and surface-water cover (f) on trends in log-transformed observed richness, rarefied richness, and abundance generated from Bayesian regression models. Models were built with biodiversity, climate, and land-use trend data for 1,758 avian communities surveyed for at least 20 years within the period of 1992–2018 as part of the North American Breeding Bird Survey; however, the trend values used for the climate and land-use predictors in the models was the slope of the fitted line plus the associated standard error. Uncertainty bars represent 95% credible intervals.

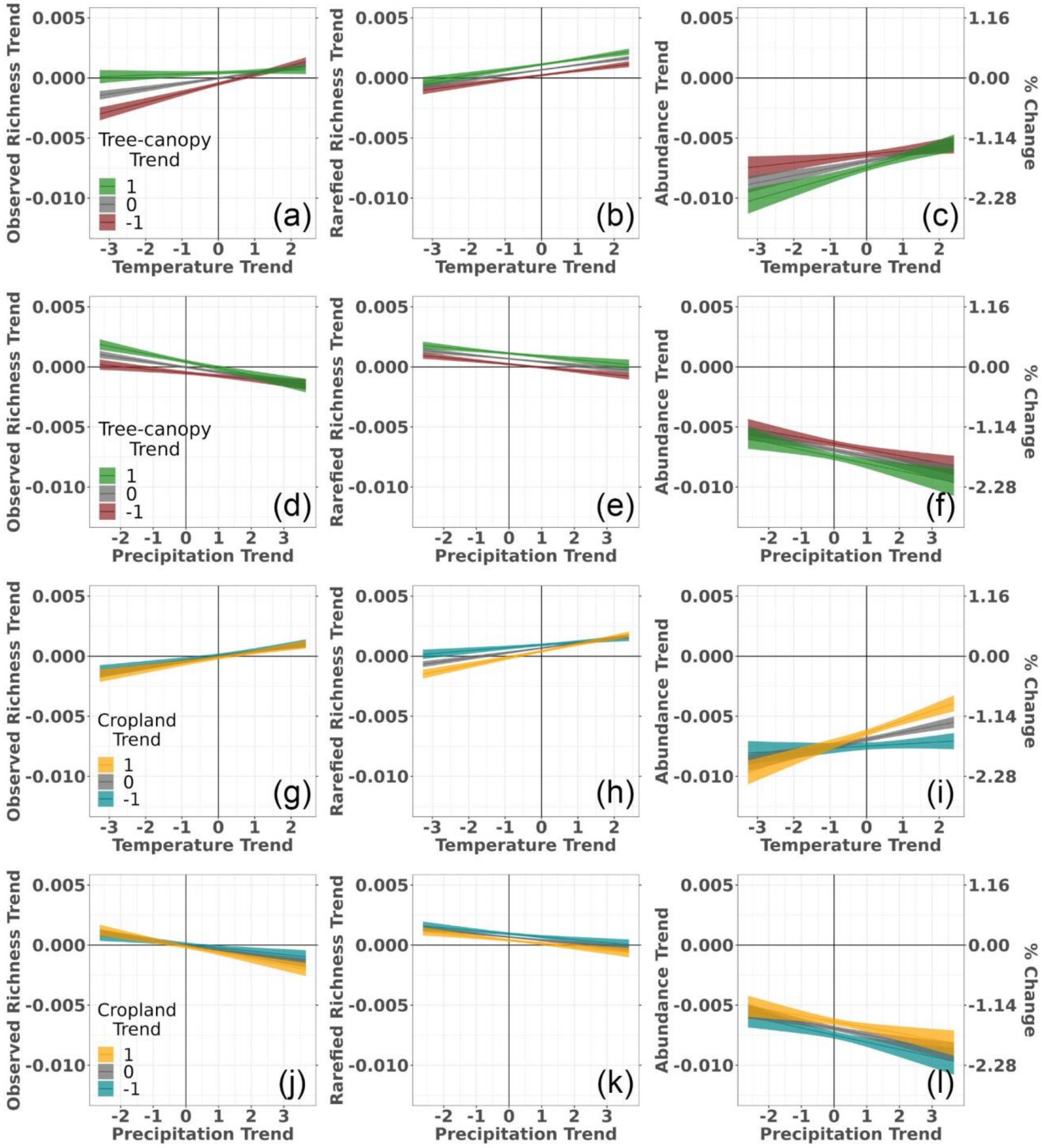


Figure S8: Upper SE of predictors model: Conditional effects of the interactions between changes in temperature and tree-canopy cover (a–c), precipitation and tree-canopy cover (d–f), temperature and cropland (g–i), and precipitation and cropland (j–l) on biodiversity trends in log-transformed observed species richness, rarefied species richness, and abundance generated from Bayesian regression models. Models were built with biodiversity, climate, and land-use trend data for avian communities surveyed for at least 20 years within the period of 1992–2018 as part of the North American Breeding Bird Survey; however, the trend values used for the climate and land-use predictors was the slope of the fitted line plus the associated standard error. Uncertainty bars represent 95% credible intervals.

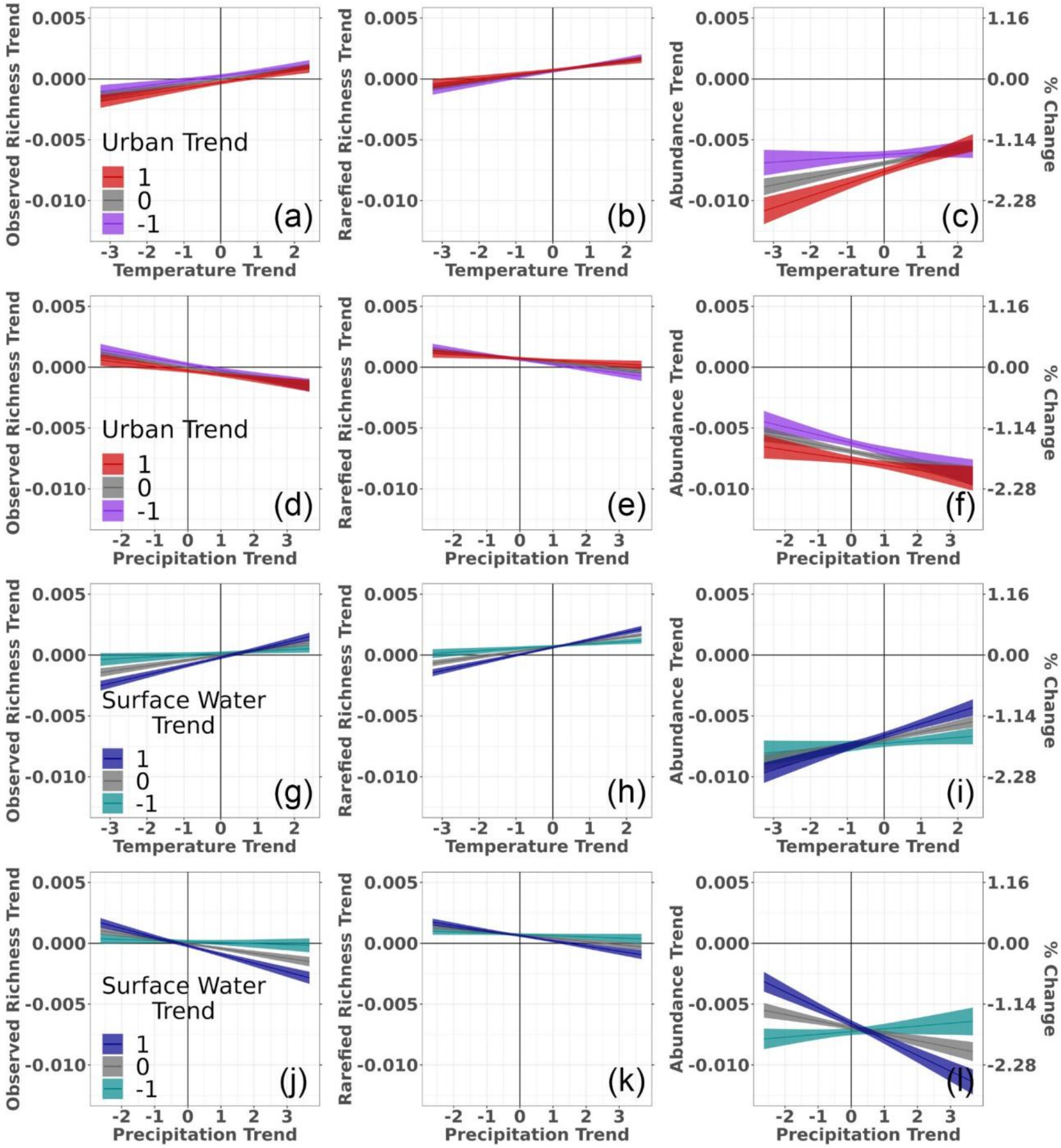


Figure S9: Upper SE of predictors model: Conditional effects of the interactions between changes in temperature and urban cover (a–c), precipitation and urban cover (d–f), temperature and surface-water cover (g–i), and precipitation and surface-water cover (j–l) on biodiversity trends in log-transformed observed species richness, rarefied species richness, and abundance generated from Bayesian regression models. Models were built with biodiversity, climate, and land-use trend data for avian communities surveyed for at least 20 years within the period of 1992–2018 as part of the North American Breeding Bird Survey; however, the trend values used for the climate and land-use predictors was the slope of the fitted line plus the associated standard error. Uncertainty bars represent 95% credible intervals.

Table S1. Comparison of the parameter estimates and credible intervals for surface-water cover change between Bayesian regression models that included the full 1.758 sample of Breeding Bird Survey routes and models that excluded 57 routes that had potentially biased trends in their surface-water cover trends. Models for each biodiversity response (log-transformed observed richness, log-transformed rarefied richness, and log-transformed abundance) are shown.

| **Model** | **Sample Size** | **Estimate** | **Lower 95% CI** | **Upper 95% CI** |
|---|---|---|---|---|
| Observed Richness | 1758 | -0.00016 | -0.00025 | -0.00006 |
| Observed Richness | 1701 | -0.00016 | -0.00026 | -0.00006 |
| Rarefied Richness | 1758 | -0.00005 | -0.00012 | 0.00002 |
| Rarefied Richness | 1701 | -0.00009 | -0.00016 | -0.00002 |
| Abundance | 1758 | 0.00035 | 0.00015 | 0.00055 |
| Abundance | 1701 | 0.00031 | 0.00011 | 0.00051 |
